# Outcome devaluation by sensory-specific satiety alters Pavlovian-conditioned behavior in male and female rats

**DOI:** 10.1101/2023.07.05.547810

**Authors:** Ankit Sood, Jocelyn M. Richard

## Abstract

Goal-directed behavior relies on accurate mental representations of the value of expected outcomes. Disruptions to this process are a central feature of several neuropsychiatric disorders, including addiction. Goal-directed behavior is most frequently studied using instrumental paradigms paired with outcome devaluation, but cue-evoked behaviors in Pavlovian settings can also be goal-directed and therefore sensitive to changes in outcome value. Emerging literature suggests that male and female rats may differ in the degree to which their Pavlovian-conditioned responses are goal-directed, but interpretation of these findings is complicated by the tendency of female and male rats to engage in distinct types of Pavlovian responses when trained with localizable cues. Here, we used outcome devaluation via sensory-specific satiety to assess the behavioral responses in male and female Long Evans rats trained to respond to an auditory CS (conditioned stimulus) in a Pavlovian-conditioning paradigm. We found that satiety-induced devaluation led to a decrease in behavioral responding to the reward-predictive CS, with males showing an effect on both port entry latency and probability and females showing an effect only on port entry probability. Overall, our results suggest that outcome devaluation affects Pavlovian-conditioned responses in both male and female rats, but that females may be less sensitive to outcome devaluation.

## Introduction

Goal-directed behavior relies on flexible mental representations of the expected value of future rewards or outcomes (Balleine & Dickinson, 1998). Cues associated with these outcomes can elicit these representations to drive goal-directed behavior, but they can also drive behaviors that are maladaptive and contribute to the psychopathology of disorders like addiction (Corbit & Janak, 2016; Ersche et al., 2016; Everitt et al., 2001; Goldstein et al., 2007; Ostlund & Balleine, 2008; Takahashi et al., 2019; Vandaele & Janak, 2018). Thus, it is important to understand the neural and behavioral mechanisms subserving the relationship between reward-predictive stimuli and mental representations of expected outcome value. Outcome devaluation, which involves reducing the value of the expected outcome, has been extensively used to test the influence of expected outcome value on action selection and action performance in instrumental conditioning paradigms (Adams & Dickinson, 1981; Mark E. Bouton et al., 2021; Parkes et al., 2016; Vandaele et al., 2017). However, less is known about the effects of outcome devaluation on Pavlovian cue-elicited reward-seeking behavior and associated neural representations of expected outcome value.

The sensitivity of Pavlovian responses to reward devaluation depends on a variety of factors, including the form of the conditioned response, the extent of training, and the method of devaluation, among others. For instance, goal-tracking behaviors in response to auditory stimuli, which preclude sign-tracking behaviors, have generally been shown to be sensitive to outcome devaluation (Johnson et al., 2009; Lex & Hauber, 2010). Devaluation via taste aversion conditioning has been shown to reduce entries and time spent in a reward port during an auditory Pavlovian cue (Holland & Straub, 1979; Johnson et al., 2009; Singh et al., 2010), as has devaluation via satiation (Lex & Hauber, 2010). On the other hand, sign-tracking responses in male rats are relatively resistant to reward devaluation via either taste aversion conditioning (Derman et al., 2018; Morrison et al., 2015; Nasser et al., 2015; Smedley & Smith, 2018) or reward satiation (Keefer et al., 2020; Patitucci et al., 2016). In addition to the above-mentioned factors, sex can also strongly influence the acquisition, intensity, and flexibility of responses to Pavlovian reward cues and devaluation (Hammerslag & Gulley, 2014; Keefer et al., 2022; Kochli et al., 2020; Madayag et al., 2017; Stringfield et al., 2019). Yet the influence of sex on the sensitivity of Pavlovian responses to outcome devaluation has only recently received any attention, and most prior studies have exclusively used male rodents. Interpretations of recent studies investigating the influence of devaluation on Pavlovian responses to lever cues in male versus female rats are complicated by sex-based differences in the tendency to sign-track in these studies and relatively low levels of goal-tracking behavior in females even under valued conditions (Keefer et al., 2022; Kochli et al., 2020). We are not aware of any published studies investigating the impact of sensory-specific satiety on Pavlovian responses to an auditory reward cue in female rats, in which subjects are disaggregated by sex.

In the present study, we investigated the effects of outcome devaluation via sensory-specific satiety on Pavlovian responses in male and female Long Evans rats. We trained rats in a Pavlovian conditioning paradigm where they learned to discriminate between two auditory cues (conditioned stimulus, CS+ and CS-), with CS+ signaling reward (10% sucrose solution) delivery and CS-signaling nothing. This was followed by within-session outcome devaluation via sensory-specific satiety and assessment of behavioral responses to CS+ and CS-presentation under extinction conditions. Both male and female rats were less likely to enter the reward port during the CS+ after devaluation. Interestingly, male, but not female rats entered the port more slowly during the CS+ following devaluation. This sex difference occurred despite similar effects of devaluation on sucrose consumption after cue testing. Overall, our results indicate that sensory-specific satiety reduces Pavlovian responses to an auditory cue in both male and female rats, but devaluation may have stronger effects on male rats. This model may be useful for investigating neural and behavioral correlates of mental representations of expected outcome value, and the influence of these representations on goal-directed behavior.

## Materials and Methods

### Subjects

Long-Evans rats (n=24, 12 males and 12 females; Envigo) weighing 250-275 grams at arrival were used in this study. Rats were pair housed by sex with *ad libitum* access to food and water and were maintained on a 14-hr/10-hr light/dark cycle (lights on at 6 am; lights off at 8 pm). Animals were handled for 5 days before beginning behavioral training. All experimental procedures were approved by the Institutional Animal Care and Use Committee at the University of Minnesota and were carried out in accordance with the Guide for the Care and Use of Laboratory Animals (NIH).

### Behavioral apparatus

All behavioral procedures were carried out in sound-attenuated operant chambers (29.53 × 23.5 × 27.31 cm) equipped with a chamber light and speakers for tone and white noise (Med Associates Inc., Fairfax, VT). One wall of the chamber had a port in the center which consisted of a receptacle with a head entry detector. A syringe pump connected with plastic tubing to the receptacle was used to deliver solutions to the receptacle. All behavioral events (lights, cue presentations, and syringe pump) were controlled by the MED-PC software (Med Associates).

### Pavlovian conditioning

Before beginning training, rats were given overnight free access in their home cages to 10% sucrose (w/v in water) and 10% Maltodextrin + 0.5% NaCl (w/v in water) on separate days to habituate them to the solutions. Following pre-exposure, rats were acclimated to the behavior chambers and consumption of the 10% liquid sucrose reward from the reward port in a magazine session which involved 30 reward deliveries without any cues in a session that lasted until all 30 reward deliveries were retrieved or 2 hours had elapsed. Subsequently, rats underwent Pavlovian conditioning as described previously (Ottenheimer et al., 2019; Richard et al., 2018). Rats were trained to discriminate between two tones: white noise and siren (ramped from 4 to 8 kHz with a 400ms period), which served as the conditioned stimuli (CS+ and CS-respectively). Each session began with the illumination of the chamber light and consisted of 30 CS+ and 30 CS-presentations. The cues lasted for 10s and were presented in a randomized fashion with variable inter-trial intervals (ITI) between cues and a mean ITI of 30s. The syringe pump turned on during the last 2s of the CS+ and delivered ∼250 ul of 10% sucrose into the reward port. The CS-did not predict any reward. Rats were trained until they met the final criteria of response probability of 0.7 or more for the CS+ trials and 0.3 or less for the CS-trials for three consecutive sessions. Response probability was calculated as the number of trials of each cue type (CS+ or CS-) with a port entry during the cue divided by the total number of trials for that cue (30). Rats that did not achieve the final criteria within 18 sessions (3 females, 1 male) were excluded from further analysis and some rats (2 females, 2 males) were used for another experiment. One male rat had to be excluded from the final analysis because of technical issues with the behavior chamber during testing. The final group size was 15 rats (7 females, 8 males).

### Reward devaluation and extinction testing

Reward devaluation was achieved via sensory-specific satiety (Colwill & Rescorla, 1985; Parkes et al., 2016). On devaluation days, rats first completed a session identical to training with 30 CS+ and 30 CS-trials as described above. This was immediately followed by a devaluation phase where rats received free access to up to 20 ml of either 10% sucrose, 10% maltodextrin + 0.5% NaCl, or no solution in the chamber for 30 minutes (LeMon et al., 2019; Vandaele et al., 2017). The order of access type was counterbalanced between animals. Immediately after the devaluation phase, rats were tested under extinction conditions which were identical to the conditioning session, except the session consisted of 5 CS+ and 5 CS-trials, and no reward was delivered with the CS+. Following the cue presentations under extinction conditions, all rats received free access to 20 ml of 10% sucrose in the chambers to assess the persistence of sensory-specific satiety. Each rat underwent 3 days of devaluation and testing such that by the end of the third session, each rat had been exposed to all the access types (10% sucrose, 10% maltodextrin + 0.5% NaCl, and no solution) once. A one-day gap, during which the rats did not perform the Pavlovian task, was given between the devaluation days.

### Behavioral data and statistical analysis

To measure the rats’ performance during Pavlovian training, we calculated the response probability, latency to enter the port, and the time spent in the port in response to the cues. A response was defined as a port entry during the 10 seconds of cue presentation. Response probability was calculated as the number of trials of each cue type (CS+ or CS-) with a port entry during the cue divided by the total number of trials for that cue (30). The latency to enter the port was measured from the start of a cue with the maximum value being 10 seconds (the end of the cue).

For the sensory-specific satiety phase where animals got free access to different solutions, the volume of solution consumed was measured and recorded. We also measured time spent in the port during the 30-minute session and divided it into 5-minute bins to assess consumption behavior over time. For the session where animals were tested under extinction conditions, the latency to enter the port and the response ratio were calculated as described above. We assessed the latency to enter the port on the first CS+ (and CS-) trial separately as this was the very first trial following the free access and without actual reward delivery and thus allowed us to assess the immediate effects of sensory-specific satiety on cue responding. Post extinction sucrose consumption and time spent in the port were measured similarly to the free access phase. Behavioral data was analyzed using linear mixed-effects models (LME) in MATLAB (Mathworks), and R using lmerTest and emmeans packages with statistical significance set at p < 0.05. Assessment of Pavlovian conditioning included fixed effects of day, cue type, and sex, with a random effect for subject. Assessment of cue responses following the devaluation manipulation included fixed effects of cue type and sex with a random effect for subject. Port entry latency following devaluation was analyzed including fixed effects of access type, sex, and trial with a random effect for subject. Analysis of time spent in port during the free access and post-extinction consumption included fixed effects of access type, sex, and time bin along with a random effect for subject. To assess if there were any potential sex differences in the behavioral performance, sex was included as a fixed effect in all LME analyses, and the data were also analyzed split by sex.

### Results

### Acquisition of discrimination between CS+ and CS-during Pavlovian conditioning

During Pavlovian conditioning, rats learned to discriminate between two auditory cues-CS+ and CS-, where CS+ predicted the delivery of the reward (10% liquid sucrose) and CS-did not predict anything (Fig. 1A-I). We quantified cue learning by calculating their probability of entering the port in response to the cues, the latency to enter the port in response to the cues, and the time they spent in the port during the cue presentation. Port entry probability during the CS+ increased as training progressed, while response probability during the CS-remained unchanged (Fig. 1A; main effect of cue type: F_(1, 928)_ = 5.102, p = 0.024; main effect of day: F_(1, 928)_ = 64.47, p< 0.001; interaction of day and cue type: F_(1, 928)_ = 29.57, p< 0.001). While we found no main effect of sex (F_(1,928)_ = 0.21, p= 0.64), we noted a significant interaction between sex and day (F_(1, 928)_ = 5.62, p= 0.01), and cue type and day (F_(1, 928)_ = 29.57, p< 0.001), but not between sex, cue type, and day (F_(1,928)_ = 2.65, p= 0.10). Therefore, we then assessed port entry probability with the data split by sex and found that both females and males displayed similar cue discrimination learning. LME analysis revealed a main effect of both cue type and day for males (Fig 1B; cue type: F_(1, 464)_ = 11.11, p< 0.001; day: F_(1, 464)_ = 197.508, p< 0.001) and females (Fig 1C; cue type: F_(1, 464)_ = 3.79, p = 0.051; day: F_(1, 464)_ = 47.98, p< 0.001). We also noted a significant interaction between cue type and day for both females (F_(1, 464)_ = 22, p< 0.001) and males (F_(1, 464)_ = 91.31, p< 0.001).

**Figure 1.**
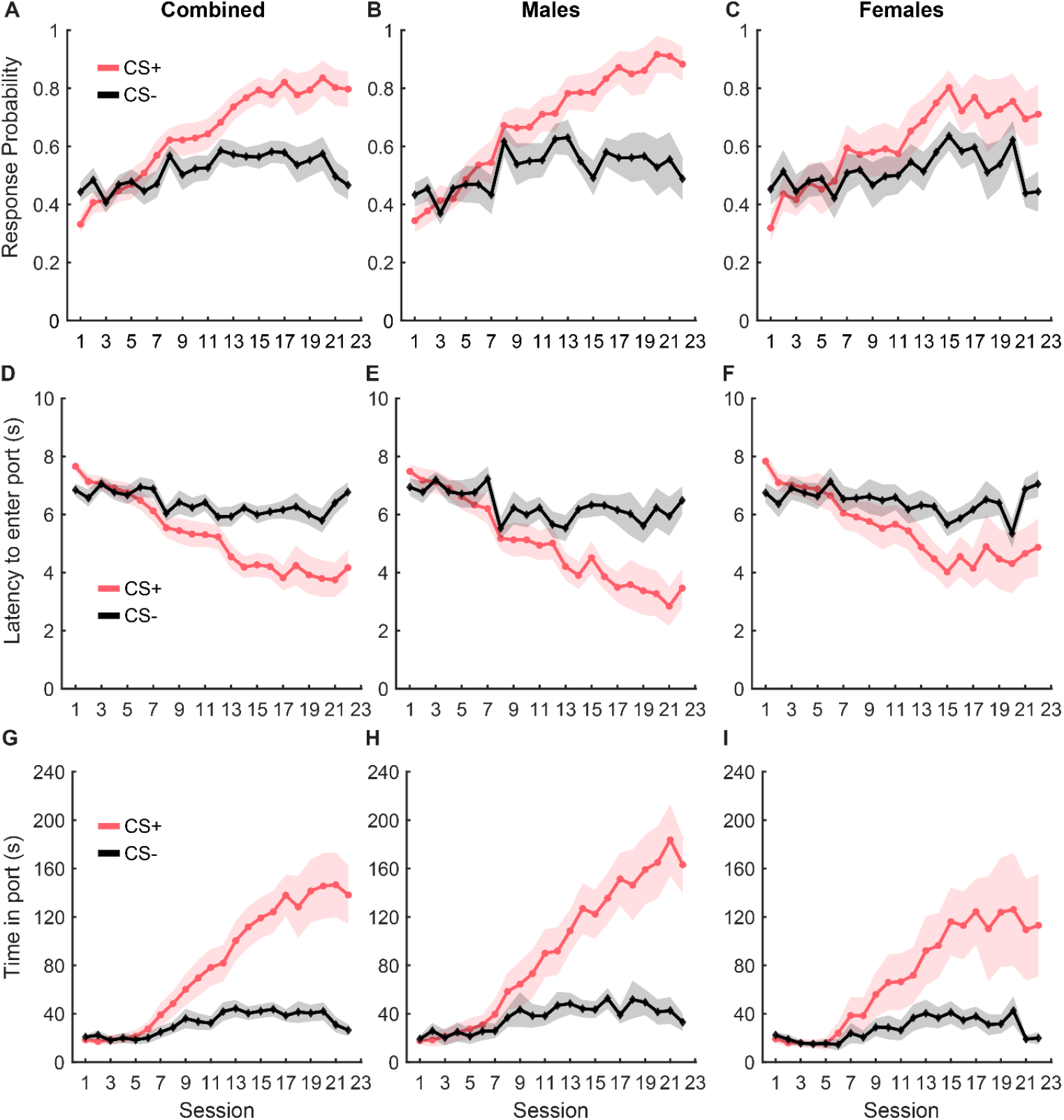
Pavlovian conditioning. The probability of entry into the reward port increased during the CS+ (red) relative to the CS-control cue (black) across conditioning in the combined group (A, n = 19), as well as separately in males (B, n = 10) and females (C, n = 9). Port entry latency during the cues in all rats (D), males (E) and females (F). Time in the reward port during the cues in all rats (G), males (H), and females (I). Data are presented as mean ± SEM.

Port entry latency provided similar evidence of cue discrimination learning. Rats decreased their latency to enter the port in response to the CS+ over training while latency in response to CS-remained unchanged (Fig 1D; main effect of cue type: F_(1, 928)_ = 7.13 p = 0.007; main effect of day: F_(1, 928)_ = 85.43, p< 0.001; interaction of cue type and day, F_(1, 928)_ = 43.13, p< 0.001). We observed no main effect of sex (F_(1,928)_ = 0.01, p= 0.89) or any interaction between sex and other factors (F_(1,928)_ = 0.001 to 3.24, p-values > 0.05), but examined the data disaggregated by sex based on the differences we observed for port entry probability. We observed a similar pattern in port entry latency for both females and males, including a significant main effect of cue type and day for both males (Fig 1E; cue type: F_(1, 464)_ = 10.20, p = 0.001; day: F_(1, 464)_ = 192.403, p< 0.001) and females (Fig 1F; cue type: F_(1, 464)_ = 5.58, p = 0.018; day: F_(1, 464)_ =66.88, p< 0.001). We also observed a significant interaction between cue type and day for males (F_(1, 464)_ = 91.96, p< 0.001) and females (F_(1, 464)_ = 33.76, p < 0.001).

Finally, rats also increased the time spent in the port in response to CS+ compared to CS-during training (Fig 1G; main effect of cue type: F_(1, 928)_ =6.68, p = 0.009; main effect of day: F_(1, 928)_ =147.02, p < 0.001; interaction between sex and day: F_(1, 928)_ = 76.86, p < 0.001; interaction between cue type and day: F_(1, 928)_ = 7.22, p= 0.007). We found no main effect of sex on time in port (F_(1, 928)_ = 0.38, p= 0.53) and no significant interaction between sex and cue type (F_(1, 928)_ = 0.78, p= 0.37) but a trend towards significance for interaction between sex, cue type, and day (F_(1, 928)_ = 3.63, p= 0.056). When data were split by sex, we observed a main effect of cue type and day for both males (Fig 1H; cue type: F_(1, 464)_ = 20.28, p < 0.001; day: F_(1, 464)_ = 349.26, p< 0.001), and females (Fig 1I; cue type: F_(1, 464)_ =5.24, p = 0.022; day: F_(1, 464)_ = 115.42, p< 0.001), and a significant interaction between cue type and day for males (F_(1, 464)_ = 180.98, p < 0.001) and females (F_(1, 464)_ = 60.34, p < 0.001). Together, our results suggest that as training progressed, both female and male rats similarly learned to discriminate between CS+ and CS-as evidenced by increased responding to CS+, decreased latency to enter the port in response to the CS+, and an increase in the amount of time spent in the port during the CS+.

### Rats consume similar amounts of sucrose and maltodextrin during free-access

To achieve devaluation via sensory-specific satiety, rats received free access to either 20 ml of 10% sucrose, 20 ml of 10% maltodextrin + 0.5% NaCl as a control solution, or no solution for 30 minutes in the behavior chambers immediately following the last training session. We measured the amount of solution consumed and the time spent by the animals in the port during the 30 min session (Fig 2). There was no difference between sucrose or maltodextrin consumption and LME analysis did not reveal a significant main effect of either access type or sex or an interaction between sex and access type (Fig 2A; F_(1,26)_ = 0.001 to 3.55, p > 0.05) with both males and females displaying similar patterns of consumption (males, Fig 2B; F_(1,14)_ = 3.22, p = 0.09; females, Fig 2C; F_(1,12)_ = 0.004, p = 0.94). To examine the consumption over time, we divided the 30-minute session into 5-minute bins and analyzed the time spent in port in each bin. LME analysis revealed a significant main effect of access type (Fig 2D; F_(1,262)_ = 36.30, p< 0.001), sex (F_(1,262)_ = 12.34, p< 0.001), and bin (F_(1,262)_ = 46.51, p< 0.001) along with a significant interaction between access type and bin (F_(1,262)_ = 21.01, p< 0.001), sex and bin (F_(1,262)_ = 7.97, p= 0.005), access type and sex (F_(1,262)_ = 6.84, p= 0.009), and access type, sex, and bin (F_(1,262)_ = 5.00, p= 0.026). Analysis of the data split by sex revealed a main effect of access type, bin, and an interaction between access type and bin for both males (Fig 2E; access type: F_(1,140)_ = 88.51, p< 0.001, bin: F_(1,140)_ = 109.51, p< 0.001, interaction: F_(1,140)_ =56.10, p< 0.001) and females (Fig 2F; access type: F_(1,122)_ = 58.65, p< 0.001, bin: F_(1,122)_ = 75.14, p< 0.001, interaction: F_(1,122)_ = 33.95, p< 0.001). These effects were likely driven by the difference between the no-solution access type and the other conditions. Rats consumed most of the solution during the first 15 minutes of the session with the time spent in port decreasing with each successive bin starting with the first. Rats spent very little time in the port after the first 15 minutes, suggesting that satiety had been achieved as a result of consumption during the first 15 minutes. This pattern was similar for both sucrose and maltodextrin across sexes and animals rarely spent time in the port on days when they did not receive access to any solution. Together, these results suggest that rats voluntarily consumed both solutions till satiety was achieved.

**Figure 2.**
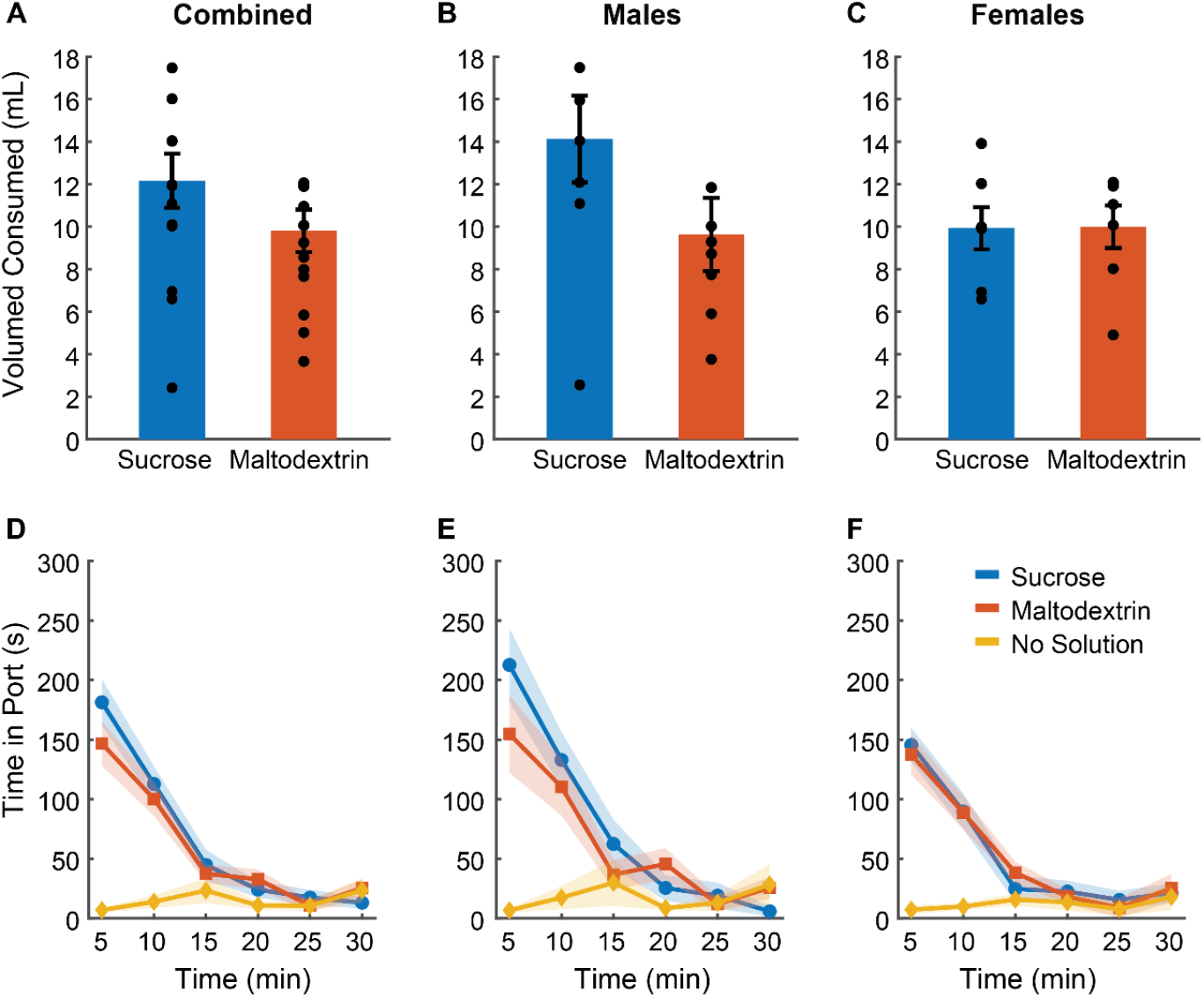
Reward devaluation via sensory-specific satiety. Free access consumption of sucrose versus maltodextrin in all rats (A, n = 19), males (B, n = 10) and females (C, n = 9). No significant differences in consumption were observed. (D) Time spent in the port during free access feeding did not differ between sucrose and maltodextrin solutions or by sex with males (E) and females (F) showing similar patterns of time in port. Data are presented as mean ± SEM, dots represent individual animals.

### Reward devaluation alters port entry latency during subsequent testing in a sex-dependent manner

To assess the effect of sensory-specific satiety on behavioral performance under extinction conditions, we analyzed port entry latency in response to CS+ and CS-immediately post the free access phase (Fig 3). Since testing under extinction conditions involved CS+ presentations without reward delivery, we analyzed latency on the first CS+ trial separately (Fig 3A-C) and across trials during the course of the session (Fig 3D-I). LME analysis of port entry latency on the first CS+ did not reveal a main effect of either access type or sex or an interaction (Fig 3A; F_(1,39)_ = 0.67 to 1.04, p > 0.05). However, when the data were split and analyzed by sex, we observed a significant main effect of access type in males (Fig 3B; F_(1,21)_ = 6.94, p= 0.004) but not females (Fig 3C; F_(1,18)_ = 0.83, p= 0.44). Port entry latency in response to the first extinction CS+ in males was highest when the rats received free access to sucrose, followed by maltodextrin, and was the least when they had access to no solution. Latency to enter the port in response to CS-did not change based on the access type during the free access phase (Fig 3A-C).

**Figure 3.**
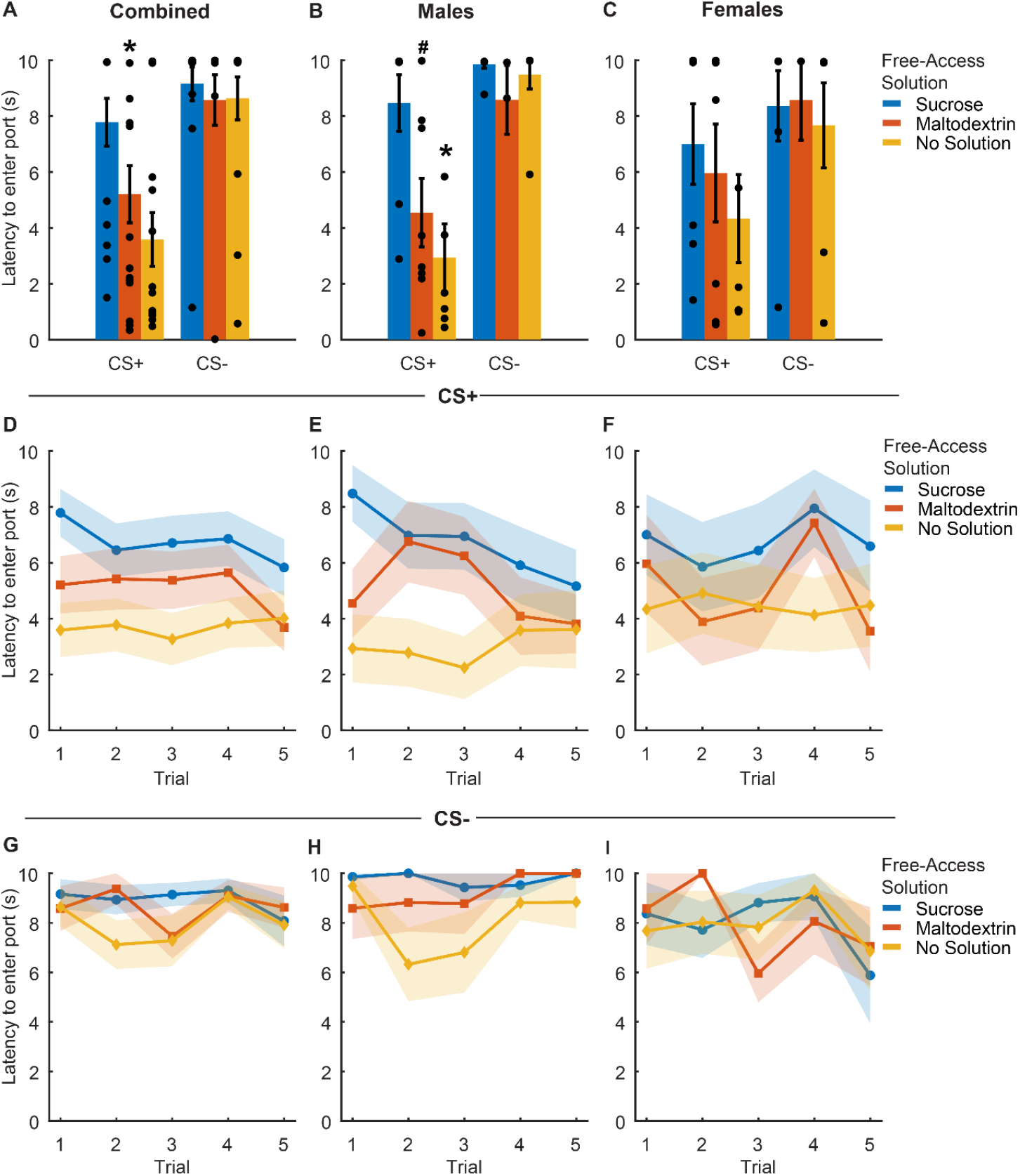
Port entry latency following devaluation via sensory specific satiety. (A-C) Port entry latency under extinction conditions in response to the first CS+ presentation versus first CS-following sucrose devaluation (sucrose free-access, blue), or control conditions (maltodextrin free-access, orange; no solution, yellow) in all rats combined (A, n = 15), males (B, n=8) and females (C, n=7). (D-F) Trial-by-trial latency to enter the port to the CS+ was longest following sucrose devaluation in all rats (D), males (E), and females (F). (G-I) Trial-by-trial latency to enter the port during the CS-was unaffected in all rats (G), males (H), and females (I). Data shown as mean ± SEM. Dots represent individual animals. *p< 0.05, #p< 0.10, compared to sucrose free-access.

We then analyzed the port entry latencies across the 5 CS+ and CS-trials to examine performance as the session progressed. LME analysis did not reveal a main effect of either access type or trial (Fig 3D; F_(1,217)_ = 0.05-0.78, p> 0.05), or an interaction between access type and trial (Fig 3D; F_(1,217)_ = 0.08, p= 0.77) for the CS+ trials. We did observe a trend towards a main effect of sex (Fig 3D; F_(1,217)_ =0.09, p= 0.07) but no significant interaction between sex and access type (Fig 3D; F_(1, 217)_ = 3.04, p= 0.08), sex and trial (F_(1, 217)_ = 2.64, p = 0.10), or sex, access type, and trial (Fig 3D; F_(1,217)_= 1.93, p= 0.16). We analyzed the data split by sex and noted a significant main effect of access type (Fig 3E; F_(1,116)_ =13.71, p< 0.001), trial (F_(1,116)_ = 5.05, p= 0.02), and a trend towards significance for interaction between access type and trial (F_(1,116)_ =3.34, p= 0.07) in males but not females (Fig 3F; access type: F_(1,101)_ =0.68, p= 0.41; trial: F_(1,101)_ =0.05, p= 0.81; access type X trial interaction: F_(1,101)_ =0.07, p= 0.78). Port entry latency across the 5 CS+ trials in males was highest when the animals had free access to sucrose, followed by maltodextrin, and the least when the animals did not receive any solution during the free access. Females showed a similar trend, but no statistical significance was noted. Latency to enter the port across 5 CS-trials remained unchanged in both males and females irrespective of the access type (Fig 3G-I). Overall, these results suggest that the access type during the free access phase affected responding to CS+ cues during testing under extinction in males, but less so in females.

### Reward devaluation via sensory-specific satiety reduces CS+ response probability

To examine whether sensory-specific satiety achieved during the free access phase led to reward devaluation, rats were tested under extinction conditions immediately following the free access phase. Rats displayed reward devaluation, as evidenced by the change in their response to CS+ depending on the solution they consumed during the free access phase (Fig 4 A-C). CS+ response probability was lowest on the day the rats received free access to sucrose and was similar to training levels on days when the rats received either maltodextrin or no solution during free access (Fig 4A; main effect of access type, F_(1,41)_ = 4.89, p= 0.03). We observed no significant effect of sex (Fig 4A; F_(1,41)_= 0.67, p= 0.41) and no interaction between sex and access type (Fig 4A; F_(1,41)_ = 0.04, p= 0.84). When the data was analyzed split by sex, LME analysis revealed a significant main effect of access type for males (Fig 4B; F_(1,22)_ = 6.74, p= 0.01) and a trend towards a significant main effect of access type for females (Fig 4C; F_(1,19)_ = 3.45, p= 0.07). Both sexes showed higher CS+ response probability on days with free access to maltodextrin or no solution compared to sucrose. Responding to CS-did not appear to change following devaluation though we observed a main effect of access type (Fig 4A; F_(1,41)_ = 9.00, p= 0.004) and an interaction between sex and access type (Fig 4A; F_(1,41)_ = 6.01, p= 0.01). This was likely due to the high response probability of males on days when they did not receive free access to any solution. When we analyzed the data split by sex, LME analysis revealed a significant main effect of access type for males (Fig 4B; F_(1,22)_ = 13.87, p= 0.001), but not females (Fig 4C; F_(1,19)_ = 2.298e-30, p =1.00).

**Figure 4.**
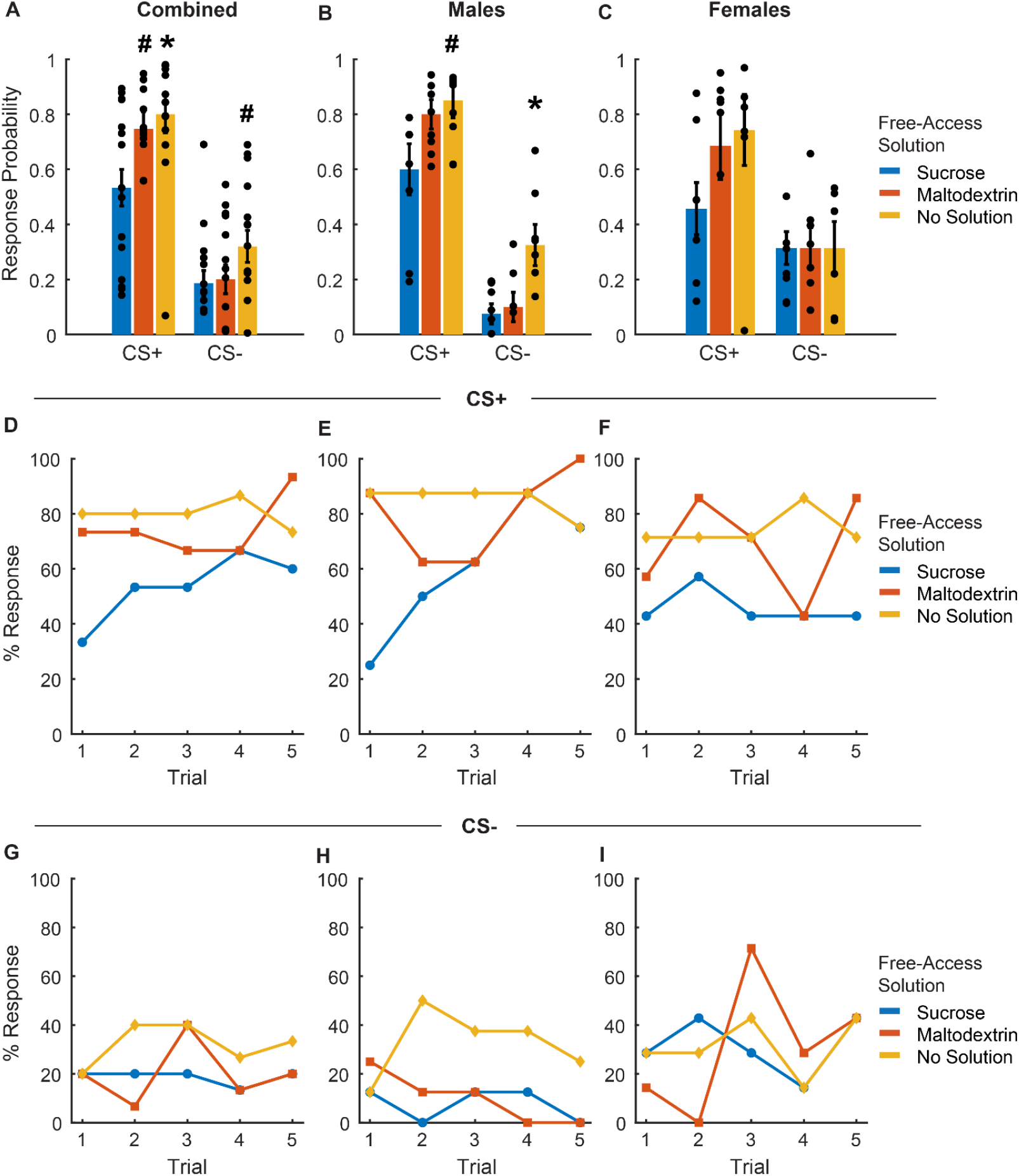
Port entry probability following devaluation via sensory specific satiety. (A-C) Port entry probability under extinction conditions in response to the CS+ versus CS-following sucrose devaluation (sucrose free-access, blue), or control conditions (maltodextrin free-access, orange; no solution, yellow) in all rats combined (A, n = 15), males (B, n=8) and females (C, n=7). Data shown as mean ± SEM. Dots represent individual animals. (D-F) Trial-by-trial responding to the CS+, shown as percentage of rats with a port entry response. The percentage of animals that responded to the CS+ with a port entry was highest following access to no solution and lowest following free access to sucrose (D). Males (E) had a higher proportion of responders on days with free access to no solution with females showing no significant difference (F) based on access type. (G-I) Trial-by-trial responding to the CS-, shown as percentage of rats with a port entry response. The proportion of animals responding to CS-did not change based on access type across sexes. *p< 0.05, #p< 0.10, compared to sucrose free-access.

To assess whether cue responding changed during the session, we analyzed the proportion of animals that responded to CS+ or CS-across trials (Fig 4D-I). LME analysis of the proportion of animals that responded on each CS+ trial did not reveal a significant main effect of either access type, sex, or trial (Fig 4D; F_(1,217)_ = 0.08 to 2.44, p> 0.05). We did observe a significant interaction between sex and trial (Fig 4D; F_(1,217)_ = 4.89, p= 0.02) and a trend toward a significant interaction between access type, sex, and trial (Fig 4D; F_(1,217)_ = 3.60, p= 0.059) but no interaction between access type and sex or access type and trial (Fig 4D; F_(1,217)_ = 0.15 to 2.59, p> 0.05). On average, the proportion of animals responding to CS+ was consistently highest for sessions when animals did not receive access to any solution, followed by sessions with access to maltodextrin, and lowest when they had free access to sucrose. When we analyzed the data split by sex, LME analysis revealed a main effect of access type (Fig 4E; F_(1,116)_ = 12.17, p< 0.05), trial (Fig 4E; F_(1,116)_ = 9.91, p< 0.05), and an interaction between access type and trial for males (Fig 4E; F_(1,116)_ = 6.50, p= 0.012) but not females (Fig 4F; access type: F_(1,101)_ = 0.57, p= 0.44; trial: F_(1,101)_ = 0.07, p= 0.78; access type X trial interaction: F_(1,101)_ = 0.13, p= 0.71). The proportion of animals that responded to CS-across trials did not differ significantly either across trials or based on the solution available during free access (Fig 4 G-I, no main effect of or interaction between access type, sex, and trial, F_(1,217)_ = 0.01 to 0.48, p-values > 0.05).

### Sensory-specific satiety affects consumption in both male and female rats

Rats displayed reduced responding to CS+ during testing under extinction conditions based on the access type they received during the free access phase. In order to further assess the effectiveness of sensory-specific satiety achieved during the free access phase, rats received free access to 20 ml of 10% sucrose, for 30 minutes in the behavior chamber immediately following the testing session. Similar to the free-access session, we measured the volume of solution consumed and the time spent by the animals in the port during the 30 min session (Fig 5). Post-extinction sucrose consumption was highest on the day when rats did not get access to any solution during the free access phase followed by the day with free access to maltodextrin, and lowest on the day with free access to sucrose (Fig 5A, main effect of access type, F_(1,41)_ = 10.85, p= 0.002). We did not observe a main effect of sex (Fig 5A, F_(1,41)_ = 0.003, p = 0.95) or an interaction between sex and access type (Fig 5A, F_(1,41)_ = 0.67, p = 0.41). We subsequently analyzed the data split by sex and LME analysis revealed a significant main effect of access type for both males (Fig 5B; F_(1,22)_ = 13.88, p = 0.001) and females (Fig 5C; F_(1,19)_ = 35.13, p< 0.001).

**Figure 5.**
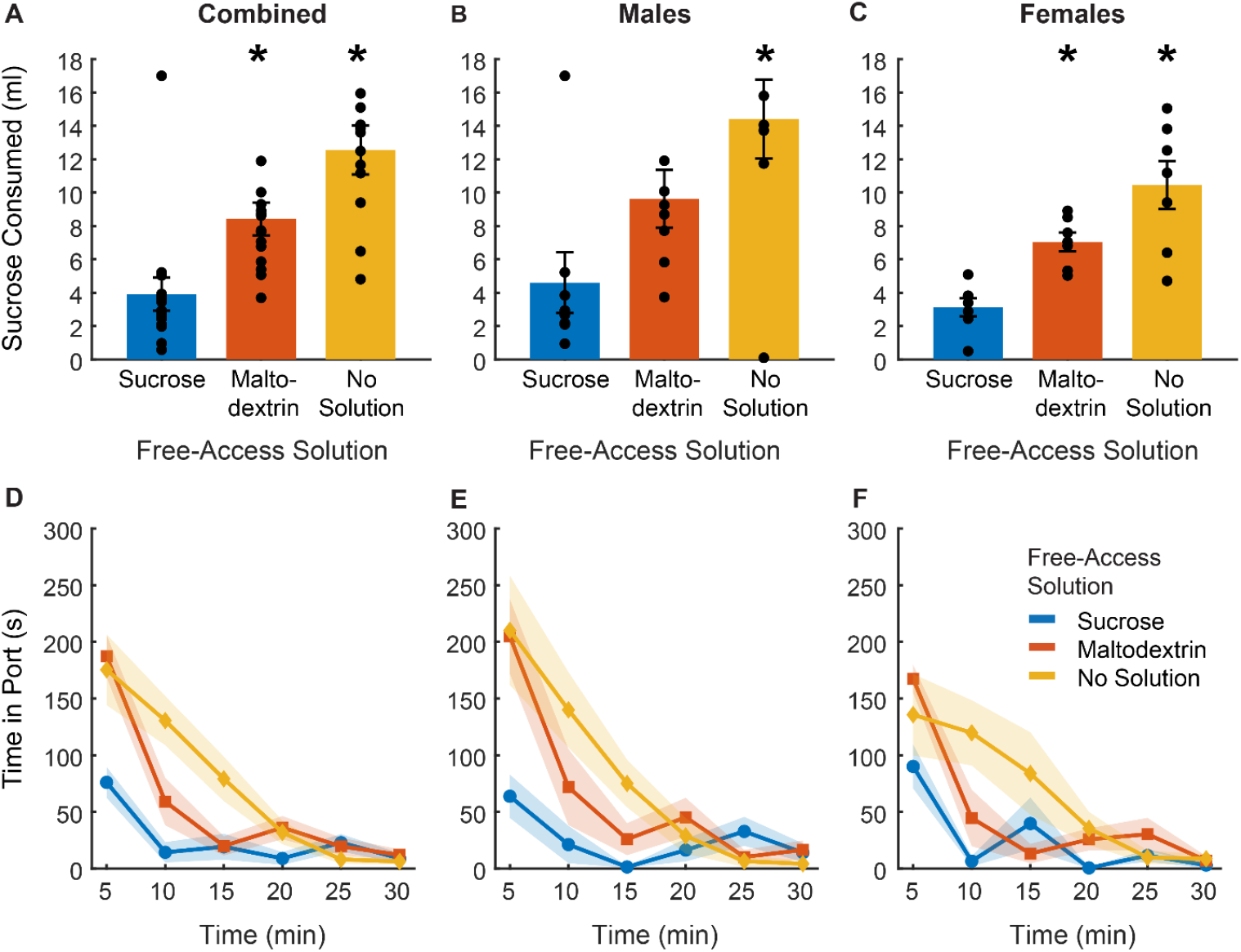
Post-testing verification of sensory-specific satiety. (A-C) Post-testing sucrose consumption was reduced by devaluation via sucrose free-access (blue) in comparison to maltodextrin free-access (orange) and no solution controls (yellow) in all rats combined (A, n =15), and similar patterns were observed in both males (B, n = 8) and females (C, n = 7). (D-F) The time spent in the port post-test was highest on days when rats had free access to no solution and least on days with free access to sucrose. Males (E) and females (F) did not differ based on the access type. *p< 0.05 compared to sucrose free-access. Data are presented as mean ± SEM. Points in A-C represent individual animals.

### LME analysis of time spent in the port during the 30-minute session revealed a similar trend

The total amount of time spent in the port was highest for the no solution access type followed by maltodextrin and least when the animals had access to sucrose during the free access (Fig 5D). LME analysis revealed a significant main effect of access type (Fig 5D; F_(1,262)_ = 11.16, p< 0.00) but not sex (Fig 5D; F_(1,262)_ = 2.00, p= 0.15), or bin (Fig 5D; F_(1,262)_ = 0.59, p= 0.44), and a significant interaction between access type and bin (Fig 5D; F_(1,262)_ = 4.6362, p= 0.03), access type and sex (Fig 5D; F_(1,262)_ = 4.6395, p= 0.03), and sex, access type, and bin (Fig 5D; F_(1,262)_ = 4.20, p= 0.04). Time spent in the port was similar for the first 5 minutes on days with free access to either no solution or maltodextrin but declined sharply for the maltodextrin access type compared to no solution over the course of the session. Rats spent significantly less time in the port from the beginning of the session on days with free access to sucrose compared with no solution and maltodextrin access days. We then analyzed the data split by sex but did not observe any sexual dimorphism in any of the measures described above. Both females and males displayed similar patterns in consumption behavior (males: Fig 5E; main effect of access type, F_(1,140)_ = 38.11, p< 0.001, interaction between access type and bin, F_(1,140)_ = 23.69, p< 0.001; Females: Fig 5F; main effect of access type, F_(1,122)_ = 14.19, p< 0.01; interaction between access type and bin, F_(1,122)_ = 5.89, p= 0.01). Overall, these results indicate that access type during the free access phase influenced the rats’ post-testing sucrose consumption regardless of sex.

## Discussion

Here, we examined the effects of outcome devaluation via sensory-specific satiety on Pavlovian-conditioned responses to an auditory cue in male and female rats. We found that free access to sucrose reduced the probability and increased the latency of port entries in response to the CS+ relative to free access to a control solution (maltodextrin) or no solution. While we did not observe any significant interactions of devaluation state and sex, when we disaggregated the data by sex we found more robust effects of devaluation on CS+ responses in male rats in comparison to female rats. This potential sex difference was not due to differences in free access consumption or differences in sucrose devaluation itself, as we observed no significant effects of sex or interactions on free access consumption, or post-test devaluation confirmation sessions. Specifically, we found that access type significantly influenced response latency in males and not females, with robust changes observed in both response probability and response latency in male rats.

During Pavlovian conditioning, associations between various qualities of the US (e.g., its sensory, hedonic, and motivational properties) and the CS are formed, and presentation of the CS can trigger representations of these qualities (Cardinal et al., 2002; Delamater & Oakeshott, 2007). The reduced behavioral performance here following outcome devaluation is indicative of its effects on the mental representation of the US triggered by the CS. We confirmed previously known findings that sensory-specific satiety reduces behavioral performance by altering the expected outcome value (Johnson et al., 2009; Lex & Hauber, 2010). We observed a reduction in behavioral performance from the first extinction CS+ following free access to sucrose suggesting that this was a satiety effect and not due to new learning during extinction. However, our study design differs from other published accounts investigating the effects of outcome devaluation on Pavlovian or instrumental behavior in some respects which are discussed below.

### Sensory-specific satiety versus general motivation

We used isocaloric liquid maltodextrin and no solution as controls as opposed to the pellets or no access only controls used in many published reports (Colwill & Rescorla, 1985; Enkel et al., 2019; Panayi & Killcross, 2022). Both maltodextrin and sucrose are preferred by rats but act on different taste receptors (Nissenbaum et al., 1987; Sclafani, 2004). The maltodextrin control allowed us to assess whether changes in cue responses reflected alterations in outcome value expectations due to devaluation of the cue-paired sucrose reward specifically, or whether these changes were due to a more general alteration in motivational state following reward consumption. Since maltodextrin never entered a CS-US association and binds to a distinct taste receptor, devaluation of the maltodextrin via free access should have a limited impact on CS-evoked responding that is driven by encoding of the expected value of sucrose. The ‘no solution’ control represented a state where both the value of the US and the animal’s general motivational state remained unaltered. Interestingly, we observed that the magnitude of cue-elicited behavioral responding after free access maltodextrin consumption was consistently in between sucrose and no solution for the different measures assessed. This slight reduction in behavioral performance compared with no solution could be indicative of an effect of maltodextrin consumption on general motivation. It is also important to note that in our design, the rats finished a training session with 30 CS-US presentations just before the free access consumption phase. It has been shown that sensory-specific habituation can lead to a reduction in within-session instrumental responding (McSweeney & Murphy, 2009). It is possible that consuming the US during the training session initiates sensory-specific satiation and subsequent consumption of maltodextrin leads to a decrease in general motivation, reflected in the slightly reduced behavioral performance. Overall, we believe that the maltodextrin control, or more generally, a control substance that shares some motivational properties with the US but is also distinguishable by the animals is worth inclusion in similar designs as it allows for dissection of sensory-specific satiety versus general motivation effects on cue-elicited behavior.

### Sex differences in Pavlovian cue-elicited behavior

Sex is an important factor known to influence reward-related behavior and existing evidence suggests that there are sex differences in behavior in response to both drug and non-drug rewards. Females, in general, tend to be more responsive during both the maintenance and extinction of drug self-administration compared to males (Chaudhri et al., 2005; Kosten & Zhang, 2008; Lynch & Carroll, 2000). With regards to nondrug rewards, it has been shown that female rats display more rapid acquisition of Pavlovian approach behavior in response to an auditory CS than males, and enhanced sign-tracking responses in cue-lever paradigms (Hammerslag & Gulley, 2014; Keefer et al., 2022; Kochli et al., 2020; Madayag et al., 2017; Pitchers et al., 2015; Stringfield et al., 2019). Females also tend to be more resistant to the effects of reward devaluation, but interpretation of this prior work has been complicated by the fact that sign-tracking responses, which tend to be more prevalent in females, are generally more resistant to devaluation than goal-tracking responses (Hammerslag & Gulley, 2014; Keefer et al., 2022; Kochli et al., 2020; Morrison et al., 2015; Nasser et al., 2015; Smedley & Smith, 2018). While we did not observe any significant sex differences in the acquisition of the Pavlovian conditioning or CS+ response probability during extinction testing, when the data were disaggregated by sex we only observed significant effects of access type in males, and not females. This suggests that females may be more resistant to outcome devaluation effects on Pavlovian responses in general, not just when those responses take the form of sign-tracking. This sex difference may be driven in part by the extensive conditioning the rats underwent prior to devaluation testing (20 sessions, 30 CS+ trials per session), which is more extensive than what is typically used for similar experiments. We predict that with less extensive training female rats may be more sensitive to outcome devaluation in this task. Overall, our observation supports the idea that female rats are more prone to displaying devaluation-resistant or habit-like behaviors and further emphasizes the importance of including both sexes in experimental designs when investigating reward-related behaviors.

## Conclusions

Here we report that auditory-conditioned Pavlovian responses are sensitive to reward devaluation via sensory-specific satiety. Rats trained to respond to a tone-CS+ signaling a sucrose reward showed reduced behavioral performance immediately following free-access sucrose consumption, indicating the sensory-specific nature of the devaluation. We observed a slight decrease in behavioral responding following free-access maltodextrin consumption, which could be due to satiety-induced effects on general motivation. Reward devaluation has been extensively used to investigate the nature of internal representations involved in the associative learning process. The most common methods of outcome devaluation are conditioned taste aversion (CTA), which involves pairing the outcome with a lithium chloride induced state of illness, and sensory-specific satiety. We chose sensory-specific satiety over CTA as the devaluation method because CTA tends to induce semi-permanent devaluation whereas satiety-induced devaluation is temporary and reversible. Additionally, the strength of the devaluation effect is influenced by the context in which devaluation is performed and satiety-induced devaluation allowed us to induce devaluation in the same context (the operant chambers) without disturbing the animals (Bouton et al., 2021; Parkes et al., 2016). As a result, our experimental design is amenable to coupling with *in vivo* approaches that aim to explore the circuit mechanisms underlying devaluation-mediated changes in behavior including *in vivo* electrophysiological recording, fiber photometry, and single-cell imaging. Since in our design all phases occur in immediate succession (training, satiety-induced devaluation, testing under extinction, and post-extinction consumption test) and without disturbing the animals, any of the above-mentioned approaches can be used to track and analyze activity changes in the same set of neurons across all phases.

## Acknowledgements

This work was supported in part by National Institutes of Health grant R01DA053208 to JMR and funding from the Medical Discovery Team on Addiction. We would like to thank Ranjani Hariharan for her assistance with behavioral training.

